# Accurate prediction of RNA secondary structure including pseudoknots through solving minimum-cost flow with learned potentials

**DOI:** 10.1101/2022.09.19.508461

**Authors:** Tiansu Gong, Fusong Ju, Dongbo Bu

## Abstract

Pseudoknots are key structure motifs of RNA and pseudoknotted RNAs play important roles in a variety of biological processes. Here, we present KnotFold, an accurate approach to the prediction of RNA secondary structure including pseudoknots. The key elements of Knot-Fold include a learned potential function and a minimum-cost flow algorithm to find the secondary structure with the lowest potential. KnotFold learns the potential from the RNAs with known structures using a self-attention-based neural network, thus avoiding the inaccuracy of hand-crafted energy functions. The specially-designed minimum-cost flow algorithm used by KnotFold considers all possible combinations of base pairs and selects from them the optimal combination. The algorithm breaks the restriction of nested base pairs required by the widely-used dynamic programming algorithms, thus facilitating the identification of pseudoknots. Using a total of 1605 RNAs as representatives, we demonstrate the successful application of KnotFold in predicting RNA secondary structures including pseudoknots with accuracy significantly higher than the state-of-the-art approaches. We anticipate that KnotFold, with its superior accuracy, will greatly facilitate the understanding of RNA structures and functionalities.

## 1 Introduction

Ribonucleic acid (RNA) are polymer molecules with essential roles involving in a large variety of biological processes [1, 2], including transcription, translation [3], catalysis [4], gene expression regulation [5], protein synthesis [6], and degradation [7]. Most biologically active RNAs, say mRNA, tRNA, and non-coding RNAs (ncRNAs), usually fold into specific structures due to the existence of self-complementary parts. These structures, together with RNA primary sequences, largely determine the biological functions of RNAs [8]; thus, a deep understanding of RNA structures is of great significance.

RNA structures can be experimentally determined using X-ray crystallography [9], nuclear magnetic resonance [10], or cryo-electron microscopy [11]. These experimental determination technologies have achieved great progress; however, the high experimental cost usually required by these technologies [12] precludes their applications – over 24 million ncRNAs have been sequenced and collected in the RNAcentral database [13] but only a tiny fraction of them have their structures experimentally determined [14]. Compared with these experimental determination technologies, computational prediction of RNA structures purely from RNA sequences is substantially effiicient and has become a promising method for understanding RNA structures.

RNA usually form secondary structure through pairing bases with hydrogen bonds and RNA secondary structures largely can be predicted without knowledge of tertiary structure as they are much more stable and accessible in cells than their tertiary form [15, 16]. One strategy for RNA secondary structure prediction is thermodynamics, which quantifies the stability of an RNA structure using folding free energy change and then selects the lowest free energy structure as it is the most probable one in the whole structure ensemble [17–19]. Turner’s nearest-neighbor model [19], a representative of thermodynamic prediction approaches, decomposes an RNA secondary structure into a collection of nearest-neighbor loops, characterizes them using multiple free energy parameters, and sums up these parameters as the free energy of the entire secondary structure [20, 21]. The free energy parameters are determined in advance using experimental techniques, say optical melting [22], or determined statistically through analyzing known RNA structures with machine learning techniques [23–25]. The lowest free energy secondary structure can be calculated using the dynamic programming technique, which determines the optimal base pairs recursively [26].

RNA structures usually contain a special kind of structure motifs called pseudoknots, which are bipartite helical structures formed through pairing a single-stranded region inside a stem-loop structure with a complementary stretch outside [27]. Pseudoknots can function as stand-alone elements or acts as parts of complex RNA structures to stabilize them [28, 29]. Understanding pseudoknots is of significant importance as pseudoknotted RNAs participate in a wide range of biological processes, including replication, RNA processing, inactivation of toxins, and gene expression control [30–32]. Figure 1a demonstrates an example of RNA secondary structure including pseudoknots. Despite the importance of pseudoknots, accurate prediction of RNA secondary structure including pseudoknots is a great challenge, partly due to the various composition of loops and helices and the lack of sequence-specific features [33]. Theoretically, the calculation of the lowest free energy structure including pseudoknots under the nearest neighbor model is NP-hard [34]. To solve this hard problem, conventional prediction approaches make compromises through limiting pseudoknot types or even focusing on the pseudoknot-free structures only. However, even if posing several reasonable limitations on pseudoknot types, the conventional dynamic programming algorithms still need *O*(*n*^4^) ∼ *O*(*n*^6^) time for an RNA with *n* bases, thus precluding their applications for long RNA sequences [35–37]. Other approaches, such as ILM [38], HotKnots [39], FlexStem [40], ProbKnot [41], and IPknot [42, 43], circumvent this computation diffiiculty using heuristic strategies. These approaches, although very fast, usually cannot guarantee quality of the predicted secondary structures. Recently, deep learning has been applied to predict base pairing probabilities with promising results [44–46]; however, the construction of secondary structure from the base pairing probabilities remains a challenge.

**Fig. 1.**
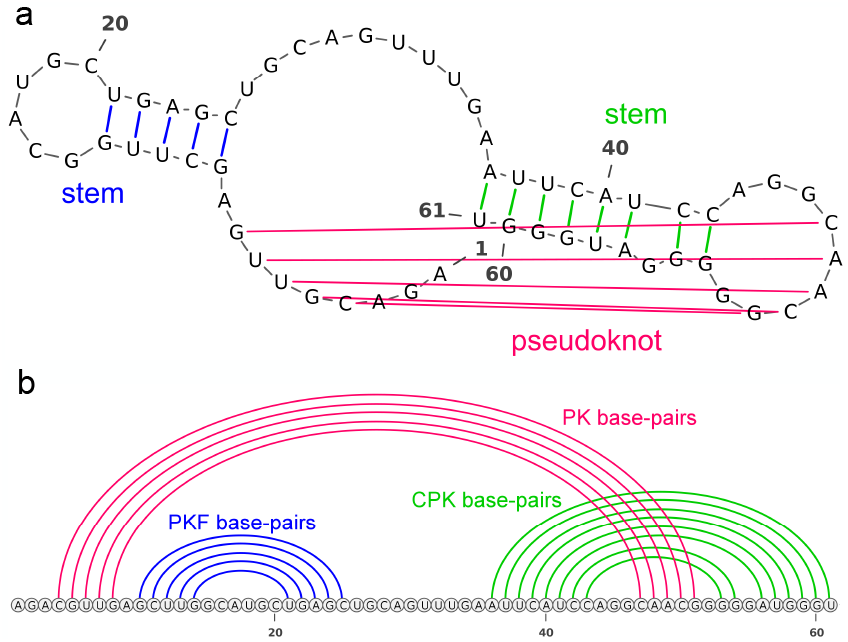
An example of RNA secondary structure including pseudoknots (bpRNA_RFAM_29722). **a**, The RNA secondary structure includes a pseudoknot formed by five base pairs: `4C-51G`, `5G-50C`, `6U-49A`, `7U-48A`, and `8G-47C`. **b**, Base pairs are divided into three categories for better evaluation of structures including pseudoknots: (*i*) pseudoknot-free (PKF) base pairs, i.e., base pairs that form no crossing with any base pair (in blue), (*ii*) pseudoknotted (PK) base pairs, i.e., the minimum set of base pairs such that, if removed, the remaining secondary structure has no pseudoknots any more (in magenta), and (*iii*) crossing-pseudoknot (CPK) base pairs, i.e., base pairs crossing some pseudoknotted base pairs (in green)

In this study, we report an accurate and fast approach (called KnotFold) to the prediction of RNA secondary structure including pseudoknots. Our approach is featured by two key elements, including: (1) *a structural potential learned using self-attention-based neural network:* KnotFold learns a structural potential from RNAs with known structures: it first predicts the base pairing probability for any two bases using a self-attention neural networks and then transforms the probabilities into a potential function. The potential function reduces the inaccuracies of hand-crafted free energies as it is learned from a large number of RNAs with known structures. Unlike the nearest-neighbor model calculating the contribution of a base pair to free energy according to its neighboring base pairs, the self-attention mechanism enables KnotFold to capture the relationship between any base pairs, especially the long-distance base pairs, thus making it more suitable for identifying pseudoknots. (2) *a specially-designed minimum-cost flow algorithm to find the secondary structure with the lowest potential:* We calculate the lowest potential structure through solving the minimum-cost flow in a flow network: the network uses nodes to represent bases and uses edges to represent base pairs with the corresponding pairwise potential as edge weight. It is worth pointing out that the minimum-cost flow algorithm considers all possible combinations of base pairs without restriction on pseudoknot types, thus making KnotFold more general and suitable for RNAs with various types of pseudoknots.

We demonstrate the accuracy of KnotFold using two benchmark sets, including PKnotTest (300 RNAs) and SPOT-TS0 (1305 RNAs). To better evaluate the performance of structure prediction approaches on pseudoknots, we divide all base pairs into pseudoknot-free (PKF) base pairs, pseudoknotted (PK) base pairs [47, 48], and crossing-pseudoknot (CPK) base pairs, derived from the convention used by previous studies including IPknot [43] (see Fig. 1b for example). For RNAs in PKnotTest, KnotFold identifies 63.1% pseudoknotted base pairs and 67.9% crossing-pseudoknot base pairs, significantly higher than the state-of-the-art approach (27.2% and 50.1%, respectively). We also provide `bpRNA_RFAM_27767` as a concrete example to investigate why the conventional dynamic programming algorithms fail. Taking `5L4O` as another example, we illustrate that KnotFold, with slight modifications, can also successfully predict base triples, which poses diffiiculty to conventional secondary structure prediction approaches. In addition, KnotFold accomplished secondary structure prediction for a long RNA with over 4300 bases within 90 seconds on an ordinary personal computer. These results clearly demonstrate the superiority of KnotFold over the existing approaches in both accuracy and effiiciency.

## 2 Results

In this section, we first demonstrate the concept of KnotFold using the RNA `bpRNA_RFAM_29722` as a representative, and then exhibit the performance of KnotFold on two datasets, including PKnotTest (containing 300 RNAs) and SPOT-TS0 [44] (containing 1305 RNAs). The details of these datasets are provided in the Methods section. We further demonstrate the advantages of KnotFold through comparing it with the existing approaches.

### 2.1 Overview of the KnotFold approach

KnotFold predicts secondary structure of a target RNA through three main steps, i.e., predicting the base pairing probability for any two bases of the given RNA, constructing a potential using the acquired base pairing probabilities, and calculating the optimal secondary structure with the lowest potential using a minimum-cost flow algorithm. We describe these steps in detail as follows.

#### Learning the base pairing probability

For an RNA sequence *x* with *n* bases, we parameterize its secondary structure as an *n × n* matrix *S* = {*S_ij_*|*S_ij_* ∈ {0, 1}, 1 ≤ *i, j* ≤ *n*}, where *S_ij_* = 1 if the *i*-th base pairs with the *j*-th base and *S_ij_* = 0 otherwise. To find the most likely secondary structure for the target RNA sequence, we first apply a deep neural network to predict the base pairing probability for any two bases. Here, we use *P* (*i* pairs with *j*|*x*) to represent the base pairing probability between the *i*-th and *j*-th bases. The neural network uses transformer encoder blocks [49] to encode bases and then calculates outer product of the encoding of two bases, which is used to represent the pairing probability of the two bases. The use of self-attention mechanism gains our approach an advantage that, when predicting the pairing probability between two bases, the entire sequence, rather than these two bases alone, is taken into consideration (see Supplementary Fig. 6 for further details of the network architecture).

#### Constructing structural potential considering all pairs of bases

To generate a secondary structure that conforms to the predicted base pairing probabilities, we construct a structural potential by summing up all pairwise potentials, i.e., the negative logarithm of the base pairing probabilities. It should be noted that, during the calculation, we correct for the over-representation of the prior by subtracting a reference distribution from the base pairing potential in the logarithm domain. The reference distribution models the base pairing probability *P* (*i* pairs with *j*|len(*x*)) independent of RNA sequence, which is computed through executing the same neural network architecture with RNA length as the only input.

In particular, the potential of a secondary structure *S* is formally described as:

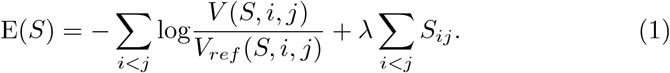

Here, *V* (*S, i, j*) is assigned the probability *P* (*i* pairs with *j*|*x*) if *S_ij_* = 1, and 1 − *P* (*i* pairs with *j*|*x*) otherwise. Similarly, *V_ref_* (*S, i, j*) represents *P* (*i* pairs with *j*|len(*x*)) if *S_ij_* = 1, and 1 − *P* (*i* pairs with *j*|len(*x*)) otherwise. The term *λ* ∑_*i<j*_ *S*_*ij*_ is introduced to penalize the inappropriate secondary structure if it has too many or too few base pairs. The parameter *λ* was optimized using the validation data set. We provide the optimal setting of this parameter in supplementary materials (Supplementary Fig. 5).

#### Calculating the optimal secondary structure

To find the optimal secondary structure *S* that minimizes the potential E(*S*), KnotFold solves a minimum-cost flow problem [50–52], in which the minimum-cost flow corresponds to the optimal secondary structure. Briefly speaking, we first constructed a bipartite graph, in which both parts consist of *n* nodes, and each node corresponds to a base of the given RNA. We drew an edge from each node in the left part to each node in the right part. We further added an extra node (called *source node*, denoted as *s*) and connected it with each node in the left part. Similarly, we also added an extra node (called *sink node*, denoted as *t*) and connected it with each node in the right part. By setting appropriate capacity and cost for each edge according to the calculated pairwise potentials, the minimum-cost flow for this network-flow problem is exactly the optimal secondary structure with the lowest potential. We used a specially-designed algorithm to solve the minimum-cost flow. The algorithm, together with the setting of capacities and costs for edges, are described in more details in Section 3.

Using the RNA `bpRNA_RFAM_29722` as a representative, we demonstrate the basic idea and main concepts of KnotFold as follows:

First, KnotFold predicted the base pairing probabilities using a deep neural network and then calculated pairwise potentials accordingly. As shown in Figure 2, the pairwise potentials exhibit three strips with significantly low values. These strips, which are perpendicular with the main diagonal, provide strong signals of three possible base pair stackings formed by the base pairing between the regions [4, 8] and [47, 51], [10, 15] and [20, 25], and [36, 43] and [53, 61], respectively. KnotFold further constructed a flow network with associated cost and capacity on edges. For example, the edge `5G-50C` and `41U-56A` are assigned with a negative cost of -8.84 and -0.47, respectively. In contrast, the edges `5G-41U`, `41U-50C` have a positive cost of 11.93 and 11.93, respectively. We assigned each edge with a capacity of 1, thus allowing any base to pair with at most one base.

**Fig. 2.**
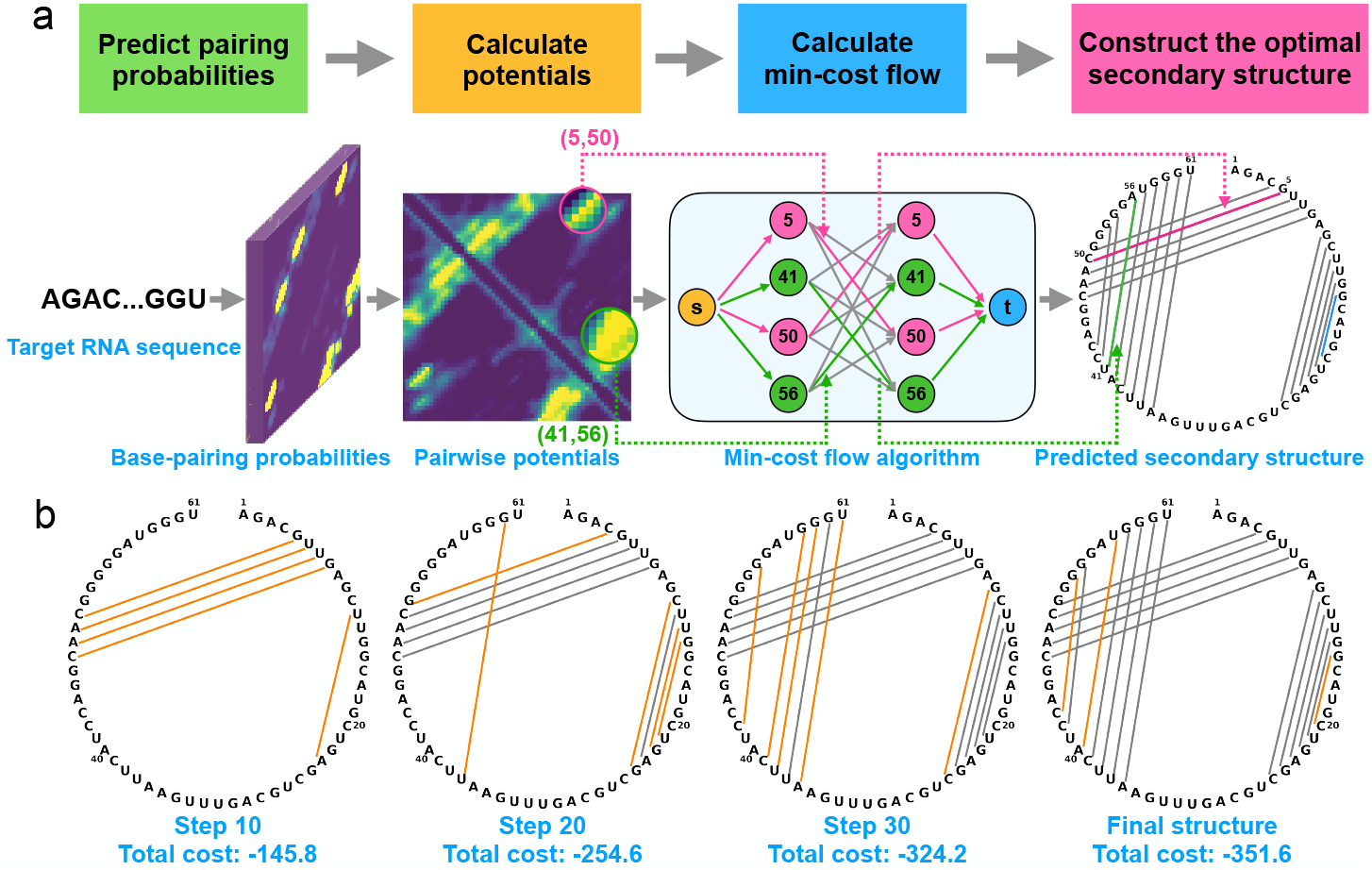
Overview of the KnotFold approach to predicting RNA secondary structure including pseudoknots. **a**, The main procedures of KnotFold illustrated using `bpRNA_RFAM_29722` as an example: KnotFold first predicts the base pairing probability for any two bases of the target RNA, then constructs pairwise potentials based on the acquired base pairing probabilities, and finally calculates the optimal secondary structure with the lowest potential using the minimum-cost flow algorithm. Here, the flow network shows four bases, i.e., `5G, 41U, 50C, 56A`, and 12 edges among these bases as representatives, and KnotFold selects the corresponding base pairs `5G-50C` (in magenta) and `41U-56A` (in green) as part of the predicted secondary structure. The final prediction consists of a total of 18 base pairs but only one false-positive base pair `15G-20C` (in blue). **b**, The iteration steps of solving the minimum-cost flow. The minmimum-cost flow algorithm begins with a zero flow with none edges and iteratively adds new edges to the current flow, or sometimes removes existing edges. We use KnotFold to construct the secondary structures corresponds to the intermediate flows. The cost decreases as iteration proceeds and finally reaches -351.6 after 36 steps. During this process, some base pairs are newly added (shown as orange lines here) while some are removed, which is described in more details in Supplementary Figure 4

Next, KnotFold calculated the minimum-cost flow using a modified shortest-path algorithm. A flow contains several paths from the source *s* to the sink *t*, and the accumulated cost of all edges traveled by the flow is denoted as its cost. To solve the minimum-cost flow, the algorithm begins with a zero-flow and continuously improved the current flow through adding, removing, or replacing some edges, in the hope of decreasing the total cost of the flow step by step. It should be pointed out that in our flow network, the flow value of each edge is either 0 or 1, i.e., an edge should be either saturated (flow value is 1) or empty (flow value is 0).

In the present case, after a total of 36 steps of improvement, the algorithm eventually acquired the minimum-cost flow with a total cost of -351.6, among which `5G-50C` and `56A-41U` are saturated with a flow while `5G-41U` and `50C-41U` are empty edges (Fig. 2b).

Finally, we obtained a predicted secondary structure using the edges traveled by the minimum-cost flow, i.e., selecting the saturated edges with the flow value of 1. In the present case, KnotFold reported 18 base pairs including `5G-50C` and `41U-56A`, and successfully identified the pseudoknot (Fig. 2).

### 2.2 Predicting secondary structures including pseudoknots using KnotFold

After demonstrating the main steps of KnotFold using `bpRNA_RFAM_29722` as an example, we further carried out a thorough evaluation of KnotFold on the PKnotTest dataset that contains a total of 300 pseudoknotted RNAs. To avoid the possible overlap between this test set and the training set, we have performed a filtering operation to guarantee that PKnotTest has no sequence with identity exceeding 80% over any RNA used for training.

The 300 RNAs in PKnotTest dataset contain a total of 21905 base pairs, which can be further divided into three categories, including 13968 pseudoknot-free (PKF) base pairs, 5325 pseudoknotted (PK) base pairs, and 2612 crossing-pseudoknot (CPK) base pairs. Thus, we can examine the prediction accuracy of KnotFold on these three categories of base pairs individually, which should facilitate the understanding of the performance of KnotFold in depth.

To investigate the contribution by the key elements of KnotFold, we built a variant of KnotFold that replaces the minimum-cost flow algorithm with the Zuker-style dynamic programming algorithm [26] for calculating the optimal secondary structure. Specifically, the variant (referred to as KnotFold-DP hereinafter) and the original KnotFold use the same pairwise potentials and they differ only in the way to infer the optimal secondary structure from these potentials.

Supplementary Table 1 suggests that KnotFold achieves a high prediction accuracy of 0.667 for all base pairs in PKnotTest, and identifies the pseudoknotted base pairs and crossing base pairs with prediction accuracy of 0.631 and 0.679, respectively. More specifically, KnotFold identifies 42.1% more pseudoknotted base pairs and 12.8% more crossing-pseudoknot base pairs than the variant KnotFold-DP, although these two approaches achieved comparable performance on the pseudoknot-free base pairs. This result clearly illustrates the advantage of KnotFold in predicting pseudoknotted base pairs.

We further carried out an in-depth examination on the failure cases of KnotFold-DP. As shown in Figure 3, despite that KnotFold-DP achieves a high accuracy of 0.868, it completely missed the five pseudoknotted base pairs, `36G-57U`, `37C-56G`, `42C-100G`, `43C-99G`, `44U-98A` (shown as dashed lines). The underlying reason is that the Zuker’s dynamic programming algorithm used by KnotFold-DP is recursive and therefore suitable for the nested base pairs. However, the pseudoknotted base pairs break this recursion assumption: when applying the dynamic programming algorithm on a pseudoknotted RNA, only a subset of base pairs can be identified.

**Fig. 3.**
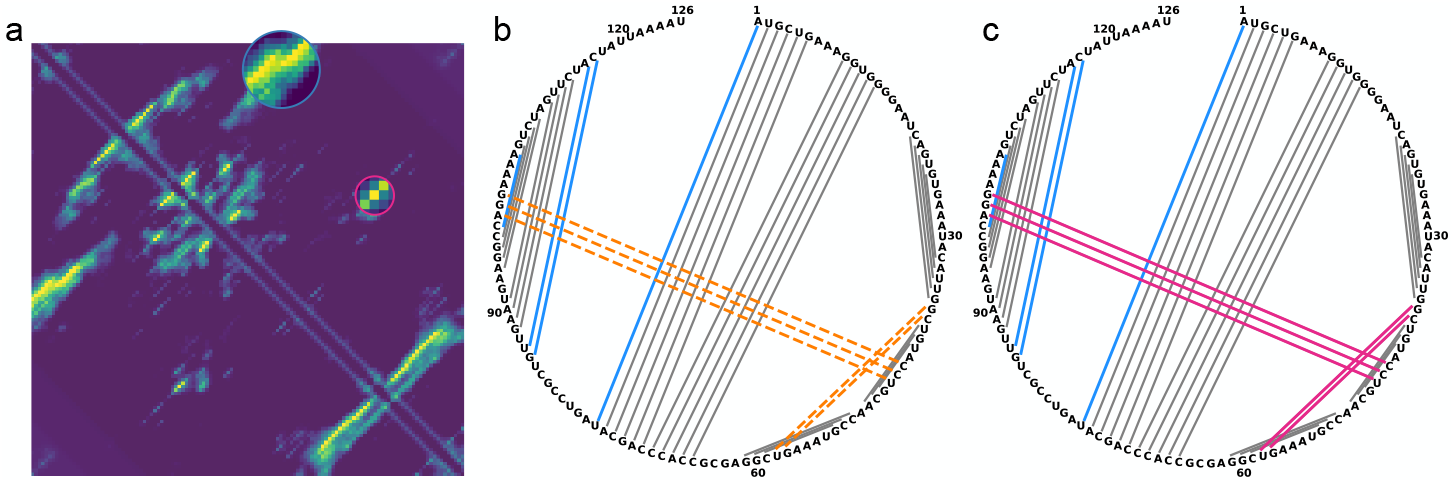
The difference between KnotFold and its variant KnotFold-DP illustrated using bpRNA_RFAM_27767 as an example. Both KnotFold and KnotFold-DP use the same pairwise potentials as their input, and they differ only in the algorithms to find the secondary structure with the lowest potential: KnotFold uses the minimum-cost flow algorithm while KnotFold-DP uses the dynamic programming algorithm. **a**, The calculated pairwise potentials for the target RNA. Here, circles highlight two regions of base pairs that are crossing. **b**, The predicted secondary structure by KnotFold-DP. The orange dash lines represent the missing base pairs while blue lines represent the false-positive base pairs. **c**, The predicted secondary structure by KnotFold. The base pairs missed by KnotFold-DP are successfully predicted (shown in magenta)

Figure 3 also suggests that KnotFold-DP correctly predicted all pseudoknot-free base pairs and 71.4% crossing-pseudoknot base pairs (solid lines) but missed the five pseudoknotted base pairs (dashed lines). In contrast, KnotFold adopts the network-flow technique and thus does not have such restrictions on the base pairs. As result, KnotFold successfully identified the five pseudoknotted base pairs.

Together, these results reveal that the major source of KnotFold’s performance comes from the use of the minimum-cost flow algorithm to identify base pairs, especially for the pseudoknotted and crossing-pseudoknot base pairs.

### 2.3 Comparison with the existing approaches

We compared KnotFold with three widely-used approaches, including RNAstructure[53], SPOT-RNA [44], and MXfold2 [54]. We provide experimental results on PKnotTest in this subsection and list the results on SPOT-TS0 in supplementary materials (see Supplementary Table 2-4).

Unlike SPOT-RNA and MXfold2 applying deep learning techniques to estimate base pairing probabilities, RNAstructure uses Turner’s nearest neighbor model to estimate free energy of an RNA structure. RNAstructure provides multiple programs to calculate the lowest free energy structure, including Fold [22], which applies the widely-known dynamic programming technique, MaxExpect [55], which reports the secondary structure with maximum expected accuracy, and ProbKnot [41], which was designed to predict secondary structure including pseudoknots. We executed different component programs to suit target RNAs: for the RNAs in SPOT-TS0, we executed all of these three programs and selected the best prediction as the final prediction of RNAstructure. In contrast, for the RNAs in PKnotTest, we directly used the prediction by ProbKnot as it was specially designed for pseudoknots.

As shown in Figure 4, KnotFold outperforms the three approaches and the superiority of KnotFold is much clearer for the crossing-pseudoknot and pseudoknotted base pairs: the accuracy of KnotFold is 0.679 and 0.631, respectively, which is considerably higher than RNAstructure (0.438 and 0.173), SPOT-RNA (0.492 and 0.272), and MXfold2 (0.501 and 0.133).

**Fig. 4.**
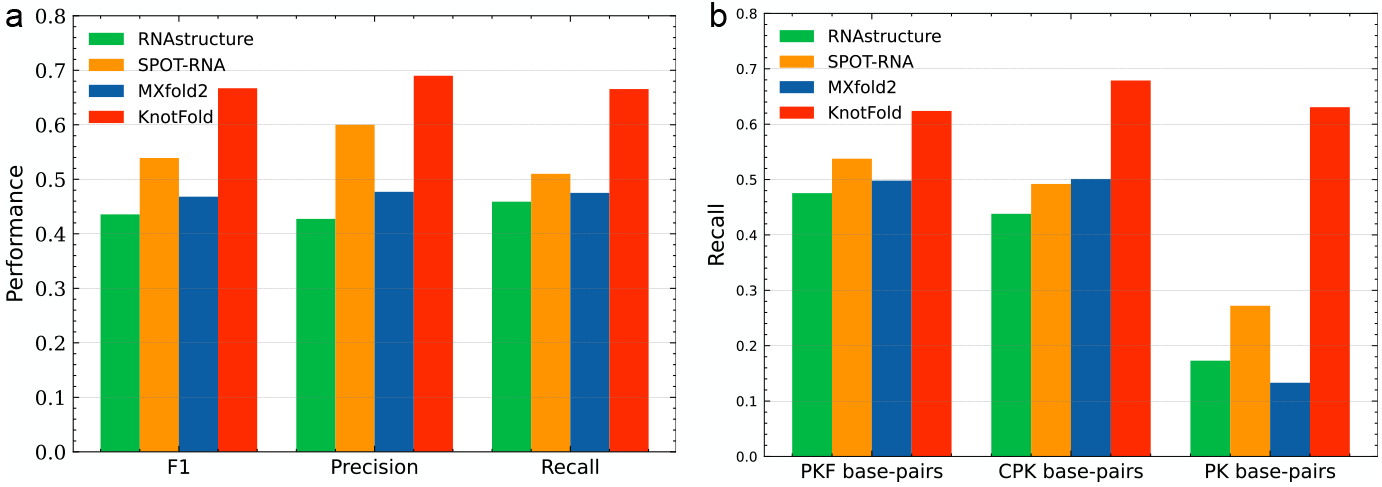
Comparison of KnotFold, RNAstructure, SPOT-RNA and MXfold2 in terms of overall prediction accuracy and the performance for various types of base pairs on PKnotTest. **a**, Overall performance (precision, recall, and F1 score) of RNAstructure, SPOT-RNA, MXfold2, and KnotFold. **b**, Recall of these approaches for various types of base pairs, i.e., pseudoknot-free base pairs, crossing-pseudoknot base pairs, and pseudoknotted base pairs

Figure 5 provides a concrete example: `bpRNA_RFAM_2518` contains five large bulges together with two pseudoknots, one connecting the regions [12, 18] and `[349, 355]`, while the other connecting the regions `[79, 82]` and `[289, 292]`. RNAstructure, SPOT-RNA and MXfold2 report secondary structures with 4, 3, and 3 bulges, respectively; however, none of them correctly identified the pseudoknots. In contrast, KnotFold successfully identified both the five large bulges and the two pseudoknots, achieving a high prediction accuracy of 0.951. We obtained a similar observation from another pseudoknotted RNA `bpRNA_tmRNA_394` (see Supplementary Fig. 1 for further details).

**Fig. 5.**
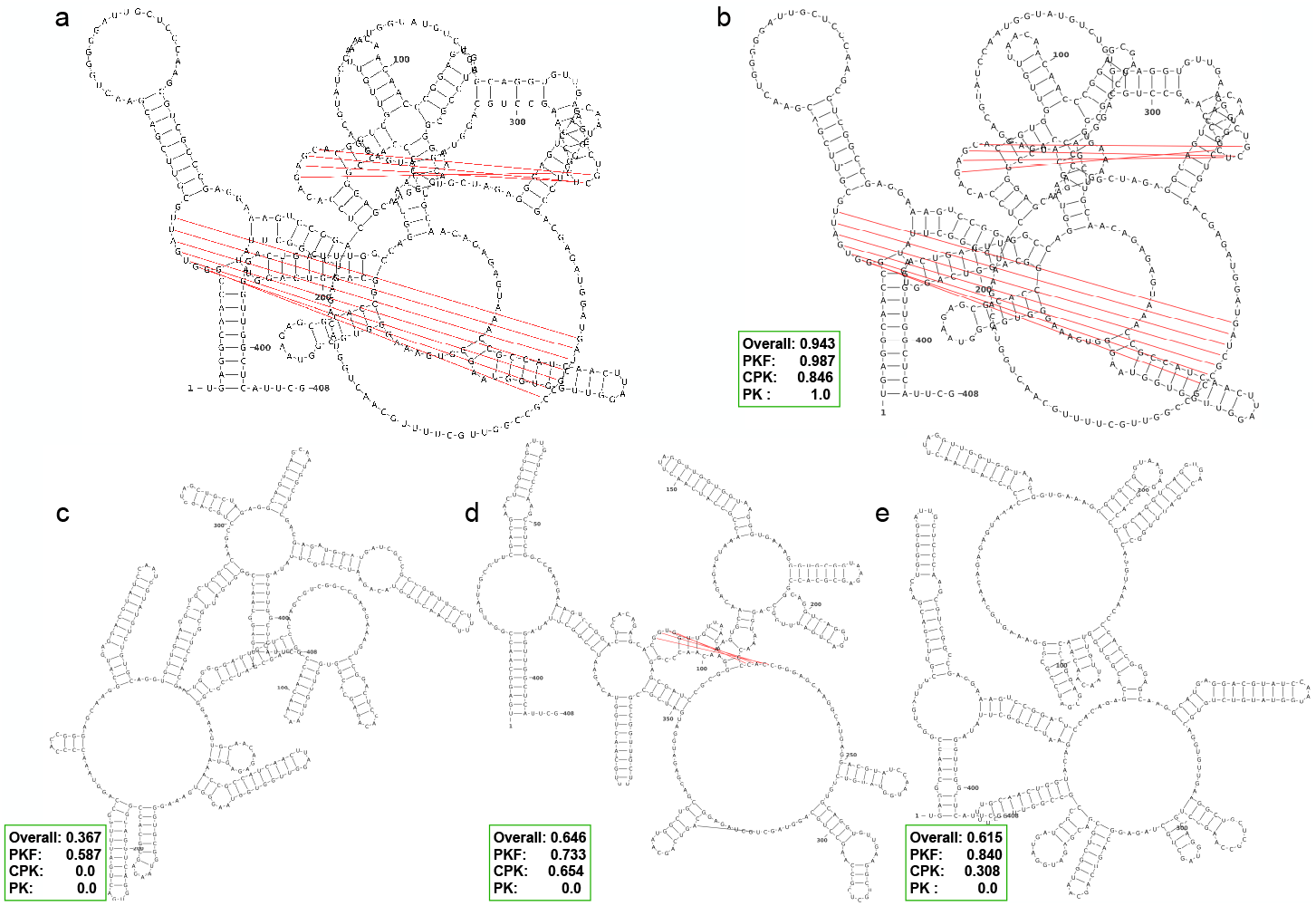
The predicted secondary structures by RNAstructure, SPOT-RNA, MXfold2 and KntoFold for bpRNA_RFAM_2518. **a**, The ground-truth secondary structure of the target RNA, in which the pseudoknotted base pairs are shown in red. The predicted structure by KnotFold (**b**), RNAstructure (**c**), SPOT-RNA (**d**) and MXfold2 (**e**) has an accuracy of 0.943, 0.367, 0.646 and 0.615, respectively. KnotFold identifies all pseudoknotted base pairs (in red) and 84.6% crossing-pseudoknot base pairs

Therefore, KnotFold shows considerable superiority in RNA secondary structure prediction, especially for pseudoknotted base pairs and crossing-pseudoknot base pairs.

### 2.4 Constructing a confidence index for secondary structure prediction

We observed a significantly tight correlation between the minimum cost reported by KnotFold and its prediction accuracy. Specifically, for 1131 RNAs in the validation dataset, the Pearson’s correlation coeffiicient between the negative average cost over saturated edges and the prediction accuracy (F1 score) is as high as 0.836 (in logarithm, Fig. 6). This tight correlation enables us to use the log value of negative average cost over saturated edges reported by KnotFold as the confidence index of prediction.

**Fig. 6.**
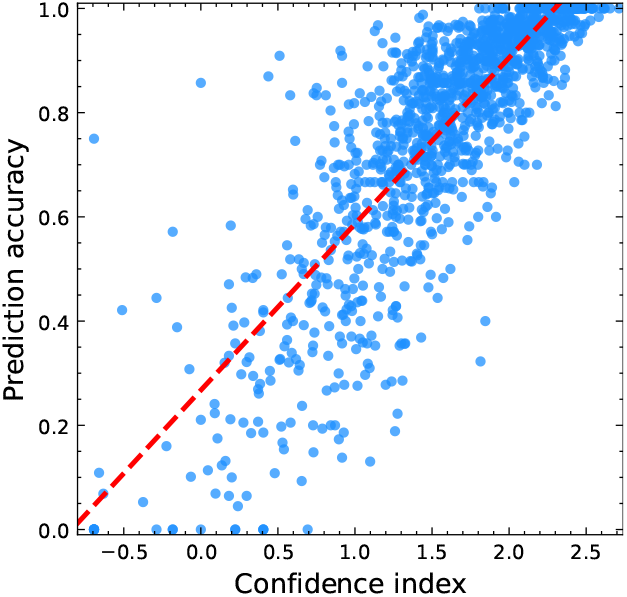
Correlation between the prediction accuracy and the estimated confidence index. We use the log value of negative average cost on saturated edges as the confidence index. For the 1131 RNAs in the validation dataset, the Pearson correlation coefficient between the prediction accuracy (F1 score) and confidence index reaches 0.836

We assessed this confidence index on the RNAs in the validation set. For example, when setting the confidence cut-off as 1.35, KnotFold reports a total of 755 RNAs, among which 717 RNAs have their prediction accuracy exceeding 0.60. This result means that, if an RNA has its confidence index estimated to be over 1.35, we can claim, with a confidence level of 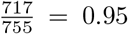, that the prediction accuracy for this RNA exceeds 0.60. We have also examined other cut-offs of the confidence index and achieved similar observations (see Supplementary Fig. 2). The construction of this confidence index will greatly facilitate the analysis of KnotFold’s prediction results and the application of the prediction approaches.

### 2.5 Extending KnotFold to identify base triples

Besides base pairs, an RNA might also form base triples [56], which involve three bases interacting edge-to-edge by hydrogen bonding. Figure 7 shows the secondary structure of the RNA with PDB entry `5L4O`, which contains three base triples, `25C-10G-45G` (in magenta), `13C-22G-46A` (in green), and `37A-29G-41C` (in blue). Previous studies have reported the importance of base triples in RNA structures and functions [57, 58].

**Fig. 7.**
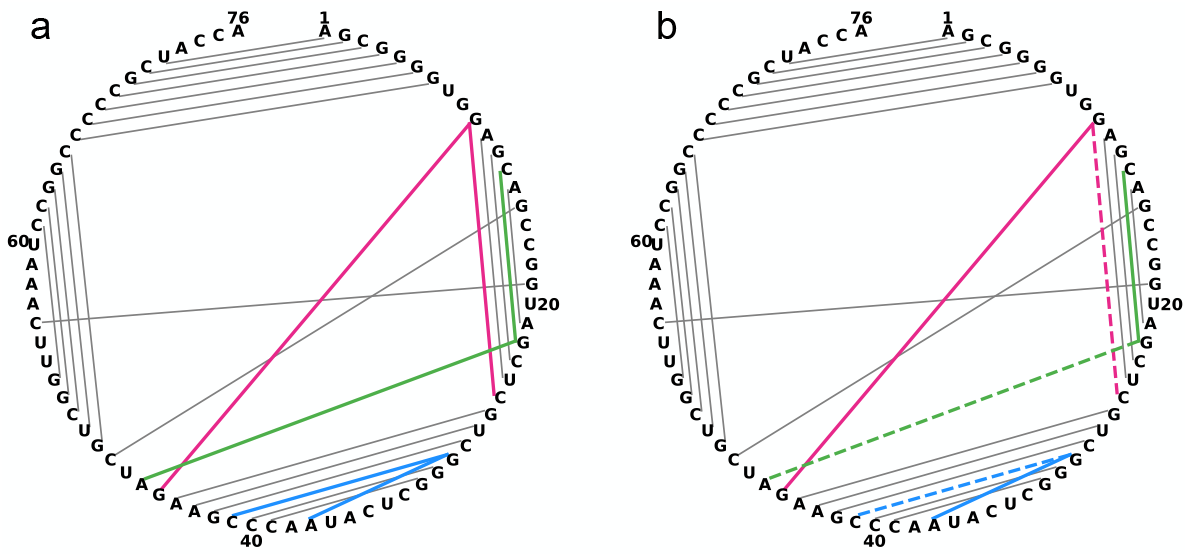
Predicting RNA secondary structure including base triples with KnotFold. **a**, The predicted structure for the RNA with PDB entry `5L4O` using the enhanced KnotFold, in which the three base triples `25C-10G-45G` (in magenta), `13C-22G-46A` (in green), and `37A-29G-41C` (in blue) are successfully identified. **b**, The predicted secondary structure by the original KnotFold without enhancement. The missing base pairs `25C-10G`, `22G-46A` and `29G-41C` (dashed lines) lead to the failure in identifying the base triples

The conventional dynamic programming algorithms, however, do not allow for base triples when constructing secondary structures. In contrast, KnotFold is capable to predict secondary structures including base triples with a slight modification without any change of its essence. In particular, we enhance KnotFold by changing the edge capacity from 1 to 2, which allows a base to interact with 2 other bases, thus forming base triples.

As shown in Figure 7, the original KnotFold missed the base pair `25C-10G` (shown as magenta dashed line), and thus failed to identify the base triple `25C-10G-45G`. Similarly, the absence of base pairs `22G-46A` and `29G-41C` leads to the failure in identifying the other two base triples `13C-22G-46A` and `37A-29G-41C`. In contrast, the enhanced KnotFold successfully identified all three base triples, and thus correctly predicted the secondary structure for this RNA. The enhanced KnotFold also successfully identified base triples for another RNA `7LYJ` (see Supplementary Fig. 3 for further details).

These results suggest that KnotFold, with slight modifications and extensions, can be used to reveal complicated motifs of RNA secondary structure, which are great challenges to the classical Zuker’s dynamic programming algorithms. This advantage will facilitate the understanding of RNA functions.

### 2.6 Effiiciency of KnotFold

Theoretical analysis suggests that for an RNA with *n* bases, KnotFold predicts RNA structure within *O*(*n*^4^) time: the prediction of base pairing probability and the subsequent calculation of pairwise potential cost *O*(*n*^2^) time, and the minimum-cost flow algorithm costs *O*(*n*^4^) time.

Despite the *O*(*n*^4^) theoretical time-complexity of KnotFold, it is extremely fast in practice: for the RNAs with as long as 2000 bases, KnotFold accomplished the entire structure prediction process within 30 seconds on an average laptop computer (Intel CPU 2.8G Hz, 16GB memory). Even for `bpRNA_CRW_55322` with 4381 bases, the longest RNA collected in bpRNA, KnotFold can accomplish its prediction within 90 seconds. These results demonstrate the high effiiciency of KnotFold and its scalability to long RNAs with even thousands of bases.

## 3 Methods

The main steps of KnotFold include predicting the base pairing probability for any two bases of the given RNA, constructing structural potential using the acquired base pairing probabilities, and calculating the optimal secondary structure with the lowest potential using the minimum-cost flow algorithm. The details of the first two steps can be referred to in the Results section and supplementary materials. In this section, we present the details of the third step as follows.

### 3.1 Transforming the secondary structure prediction problem into a minimum-cost flow problem

As described above, we predict the secondary structure for a target RNA through finding the secondary structure with the lowest potential. The potential function, which is shown in Equation 1, is the accumulated pairwise potential of all possible pairs of bases. The key insight of our approach is that, although the potential is defined as the sum of pairwise potentials of all possible pairs of bases, it is essentially determined by the base pairs appearing in the secondary structure only. Specifically, we defined a novel measure, called *cost*, for each pair of bases and proved that the potential can be rewritten as the total cost of the base pairs that form secondary structure, i.e.,

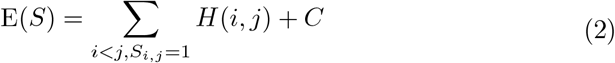

Here, *C* represents a constant number independent of the given RNA, and *H*(*i, j*) represents the cost of the base pair (*i, j*) and is described as:

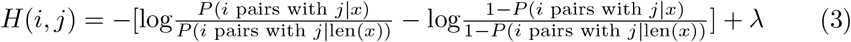

We provide the strict proof of the equivalence between Equation 1 and Equation 2 in supplementary materials.

The equivalence between Equation 1 and Equation 2 enables us to transform finding the secondary structure with the lowest potential into calculating the minimum-cost flow in an appropriately-designed network. In particular, the network consists of a bipartite, and each part of the bipartite consists of *n* nodes that represent the *n* bases of the target RNA. We connected every node in the left part to every node in the right part with an edge, which essentially represents a possible base pair. For the edge (*i, j*) connecting the *i*-th base and the *j*-th base, we set its capacity as 1 and its cost as *H*(*i, j*).

In this flow network, the base pairs represented by the minimum-cost flow essentially form a secondary structure with the lowest potential. The details of the network construction are described as follows.

#### Algorithm 1 Constructing the flow network *G*

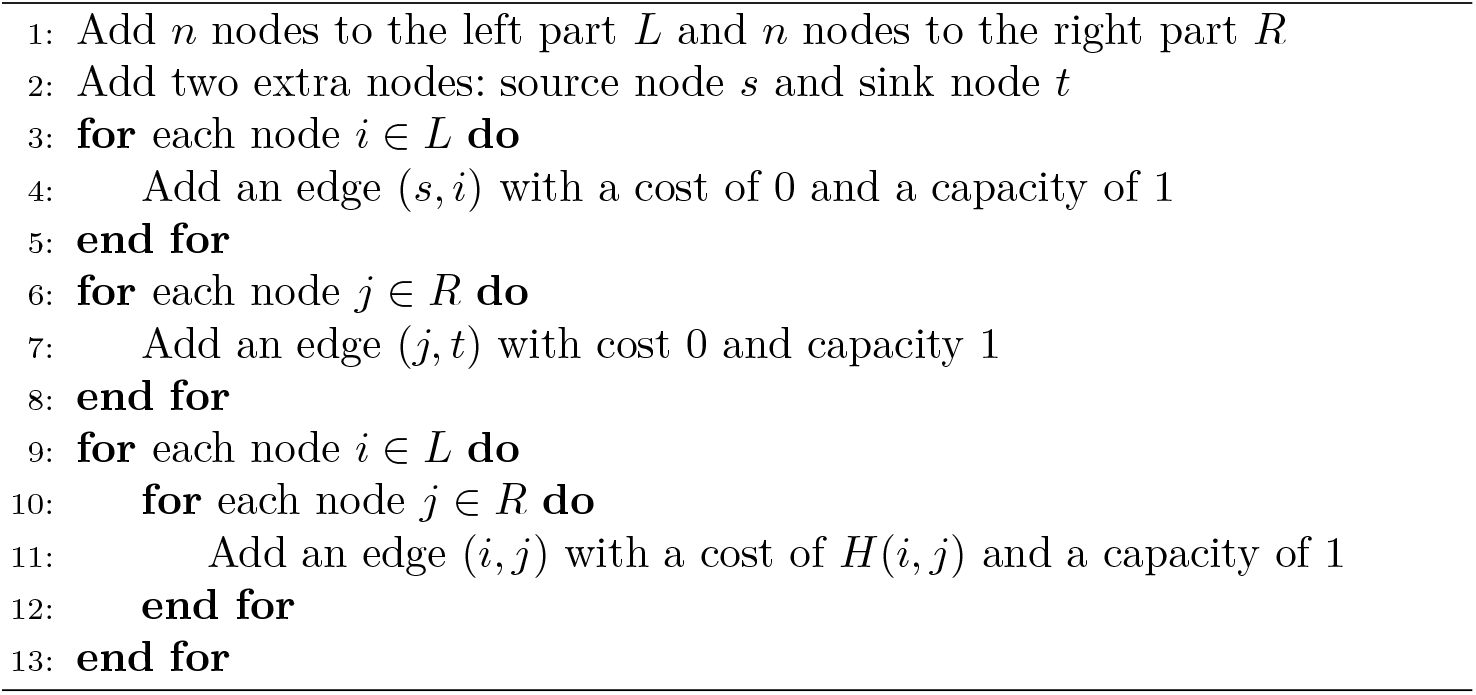

### 3.2 Solving the minimum-cost flow using a modified shortest-path algorithm

We solve the minimum-cost flow in the constructed flow network using a modified shortest-path algorithm. Specifically, we start from a 0-flow, i.e., all edges are initialized with a flow value of 0. Next, we iteratively execute the following two steps:

i. Constructing a residual graph *G_f_* according to the current flow *f*. For each edge (*i, j*) in the flow network, we add two edges in the residual graph *G_f_*, including a forward edge (*i, j*) with capacity 1 − *f* (*i, j*) and cost *H*(*i, j*), and a backward edge (*j, i*) with capacity *f* (*i, j*) and cost −*H*(*i, j*).
ii. Finding the shortest path from the source *s* to the sink *t*, denoted as *s* − *t* path, in the residual graph *G_f_*, followed by pushing along this path to augment the current flow *f*. Here, the shortest path from *s* to *t* refers to the path with the minimum accumulated cost of the edges traveled by this path.

Finally, we extract the saturated edges from the minimum-cost flow, i.e., the edges with a flow value of 1, and report a secondary structure with base pairs corresponding to these saturated edges as the predicted secondary structure. Unlike the classical shortest-path algorithm, we use a modified stopping criterion: the two steps are executed until no *s* − *t* path with positive accumulated cost can be found in the residual graph. With this stopping criterion, the modified shortest-path algorithm can solve the minimum-cost flow.

The details of the modified shortest-path algorithm are provided in Algorithm 2.

#### Algorithm 2 Modified shortest-path algorithm for minimum-cost flow

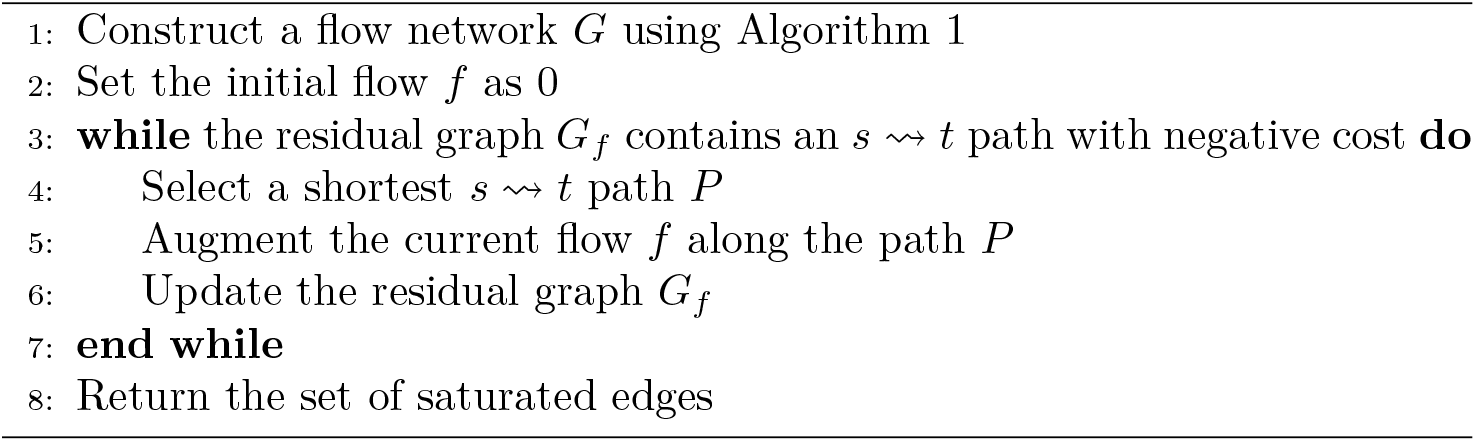

### 3.3 Dataset

In the study, we evaluate the prediction approaches using the RNAs extracted from the following three databases:

i. bpRNA-1m: one of the most comprehensive datasets of RNA secondary structures [59]. bpRNA-1m contains 102,318 sequences extracted from multiple datasets including Rfam 12.2. We utilize its RNA sequences and secondary structures for training and validation, and we build a pseudoknot test dataset PKnotTest from bpRNA-1m [59].
ii. Rfam: Rfam (version 14.7) contains RNAs covering 4069 families [60]. We use the newly added RNAs after the release of Rfam (version 12.2) to construct training and validation datasets.
iii. Protein Data Bank (PDB): We use high-resolution RNA 3D structures (<3.5 Å) collected in PDB to assess the prediction of base triples [14].

Using the RNAs collected in bpRNA-1m [59] and Rfam 14.7 [60], we prepared training set, valildation set, and test set as follows: To reduce the potential redundancy existing in these RNAs, we clustered them using CD-HIT-EST [61] at 80% sequence-identity cutoff and select only one representative RNA from each cluster. For the sake of fair comparison with the existing approaches SPOT-RNA and MXfold2, we discarded the clusters that have overlap with SPOT-TSO, which was used by the two approaches. As results, we acquired a total of 20171 non-redundant RNAs.

From these non-redundant RNAs, we randomly selected 300 RNAs, which include pseudoknots in their secondary structures, and use them as test set (denoted as PKnotTest). The remaining RNAs were randomly split into a training set and a validation set, which contain 18740 and 1131 RNAs, respectively.

### 3.4 Evaluation criteria

We evaluated the prediction accuracy using the same metrics as SPOT-RNA and MXfold2, which include precision, recall, and F1 score. We calculated the average precision, recall, and F1 score to evaluate the overall performance on a dataset and the average of recall to evaluate the performance on different types of base pairs on PKnotTest.

## 4 Conclusion

The results presented here have highlighted the special features of KnotFold: it uses a deep neural network to learn a structural potential that considers all pairs of bases, thus making it suitable for identifying long-distance base pairs, especially pseudoknots; it also uses a specially-designed minimum-cost algorithm to find the secondary structure with the lowest potential. Using a total of 1605 RNAs collected in popular benchmark datasets as representatives, we demonstrate the accuracy and effiiciency of KnotFold, together with its superiority over the existing approaches.

We also analyzed the source of of the power of KnotFold through comparing it with its variant, which combines the classical Zuker’s dynamic programming algorithm and the pairwise potentials predicted by deep neural networks. The analysis suggested that the main source of the power of KnotFold to predict pseudoknots comes from the application of a minimum-cost flow algorithm to calculate the secondary structure with the lowest potential.

The ideas of KnotFold can be readily extended without significant modifications to solve other complicated structure motifs. For example, when changing the capacities over edges from 1 to 2, KnotFold can easily solve base triples, which represents a great challenge to classical Zuker’s dynamic programming algorithms.

Although the minimum-cost flow algorithm constructs structure with the minimum potential, the accuracy of KnotFold relies heavily on the predicted probabilities of base pairing and the subsequent calculation of pairwise potentials. For example, the accuracy of base pairing probability is usually low for the long RNAs with over 2000 bases, or the rare RNAs with special secondary structure types. In this case, even if using the minimum-cost flow algorithm, the predicted secondary structures are not convincing. How to improve the prediction of base pairing probabilities is one of our future works. In addition, except for the optimal secondary structure with the lowest potential, the calculation of sub-optimal secondary structures might yield a secondary structure ensemble, which will provide a deep insight into the predicted secondary structure.

We anticipate that KnotFold, with its superiority in accuracy and effiiciency, will greatly facilitate our understanding of RNAs with complicated structures and their biological functions.

## 5 Acknowledgements

We would like to thank the National Key Research and Development Program of China (2020YFA0907000), and the National Natural Science Foundation of China (32271297, 62072435, 31770775, 31671369) for providing financial supports for this study and publication charges.

## 6 Author contributions

D.B. directed the RNA secondary structure prediction project and revised the manuscript. T.G. designed the approach, did the experiments, and drafted the manuscript. F.G. implemented the minimum-cost flow algorithm.

## 7 Competing interests

The authors declare no competing interests.

## Notes

https://github.com/gongtiansu/KnotFold

### Competing Interest Statement

The authors have declared no competing interest.

